# Infant and Adult Human Intestinal Enteroids are Morphologically and Functionally Distinct

**DOI:** 10.1101/2023.05.19.541350

**Authors:** Grace O. Adeniyi-Ipadeola, Julia D. Hankins, Amal Kambal, Xi-Lei Zeng, Ketki Patil, Victoria Poplaski, Carolyn Bomidi, Hoa Nguyen-Phuc, Sandra L. Grimm, Cristian Coarfa, Fabio Stossi, Sue E. Crawford, Sarah E. Blutt, Allison L. Speer, Mary K. Estes, Sasirekha Ramani

**Author notes:** Corresponding author: Sasirekha Ramani, Ph.D; Department of Molecular Virology and Microbiology, 1 Baylor Plaza RM 939E, Houston, TX, 77030. Julia D. Hankins: University of Kansas, Lawrence, KS.

## Abstract

**Background & Aims:** Human intestinal enteroids (HIEs) are gaining recognition as physiologically relevant models of the intestinal epithelium. While HIEs from adults are used extensively in biomedical research, few studies have used HIEs from infants. Considering the dramatic developmental changes that occur during infancy, it is important to establish models that represent infant intestinal characteristics and physiological responses.

**Methods:** We established jejunal HIEs from infant surgical samples and performed comparisons to jejunal HIEs from adults using RNA sequencing (RNA-Seq) and morphologic analyses. We validated differences in key pathways through functional studies and determined if these cultures recapitulate known features of the infant intestinal epithelium.

**Results:** RNA-Seq analysis showed significant differences in the transcriptome of infant and adult HIEs, including differences in genes and pathways associated with cell differentiation and proliferation, tissue development, lipid metabolism, innate immunity, and biological adhesion. Validating these results, we observed a higher abundance of cells expressing specific enterocyte, goblet cell and enteroendocrine cell markers in differentiated infant HIE monolayers, and greater numbers of proliferative cells in undifferentiated 3D cultures. Compared to adult HIEs, infant HIEs portray characteristics of an immature gastrointestinal epithelium including significantly shorter cell height, lower epithelial barrier integrity, and lower innate immune responses to infection with an oral poliovirus vaccine.

**Conclusions:** HIEs established from infant intestinal tissues reflect characteristics of the infant gut and are distinct from adult cultures. Our data support the use of infant HIEs as an ex-vivo model to advance studies of infant-specific diseases and drug discovery for this population.

**Importance:** Tissue or biopsy stem cell-derived human intestinal enteroids are increasingly recognized as physiologically relevant models of the human gastrointestinal epithelium. While enteroids from adults and fetal tissues have been extensively used for studying many infectious and non-infectious diseases, there are few reports on enteroids from infants. We show that infant enteroids exhibit both transcriptomic and morphological differences compared to adult cultures. They also differ in functional responses to barrier disruption and innate immune responses to infection, suggesting that infant and adult enteroids are distinct model systems. Considering the dramatic changes in body composition and physiology that begins during infancy, tools that appropriately model intestinal development and diseases are critical. Infant enteroids model key features of the infant gastrointestinal epithelium. This study is significant in establishing infant enteroids as age-appropriate models for infant intestinal physiology, infant-specific diseases and responses to pathogens.

## Introduction

The human small intestine is a highly dynamic organ that regulates digestion, nutrient absorption, and waste elimination. It also serves as a barrier against potential luminal pathogens. Infectious and non-infectious pathologies of the small intestine contribute significantly to disease burden across all ages. However, clinical presentations and the prevalence of some diseases may vary with age. For example, severe dehydrating gastroenteritis due to rotavirus occurs primarily in children under the age of 5 years although infections can occur at all ages (1). By contrast, diarrhea and vomiting due to norovirus occurs in persons of all ages (1). Necrotizing enterocolitis (NEC) almost exclusively affects neonates while age-related differences in the presentation and course of other gastrointestinal conditions such as inflammatory bowel diseases have been reported (2–4). In order to understand the pathophysiological mechanisms of disease and to design targeted preventive or therapeutic interventions, it is important to establish models that appropriately represent the developmental stage and physiology of the human small intestine across the lifespan.

The small intestinal epithelium is composed of a polarized epithelial layer that contains different cell types including enterocytes, enteroendocrine cells, tuft cells, goblet cells, Paneth cells and stem cells (5). Common models to study small intestinal disorders include transformed cell lines and animals which, while useful, do not always efficiently recapitulate the physiology and microenvironment of the human small intestine. Human intestinal enteroids (HIEs) are self-organizing three-dimensional *ex vivo* tissue cultures derived from stem cells in human intestinal surgical tissues or biopsies (5). These non-transformed cultures contain multiple intestinal cell types, and thus reflect *in vivo* epithelial heterogeneity (5–7). Once established, HIEs can be passaged long-term making them a very valuable laboratory tool (5, 6). HIEs have been widely used to interrogate gut physiology (7–12), host-pathogen interactions (13–19), drug activity (20, 21), and cell-to-cell communication (22, 23). Although HIEs do not completely reflect the complexity of the intestine in terms of the immune and stromal cells, the enteric nervous system, or the microbiome, these transformative cultures are expanding our understanding of the cellular composition, morphology, and functionality of the human intestinal epithelium.

HIEs from adult intestinal tissues or biopsies are used extensively in biomedical research and are well characterized (13–15, 18). Human fetal tissue-derived enteroids have also been used to study the pathogenesis of different infectious agents (18, 19) and for diseases such as NEC (24). Currently, there are only a limited number of studies using HIEs from infants and young children and very few systematic comparisons have been made between the different models (25, 26). Characteristics of fetal HIEs correlate with their developmental age and demonstrate immature metabolic and host-defense functions compared to adult HIEs (24, 27). Differences in transcriptional responses between adult and preterm enteroids to *Enterococcus faecalis* have been described (26). A study using HIEs from 2- and 5-year-old children showed significantly shorter enterocyte cell height and lower transepithelial electrical resistance (TEER) than adult HIEs (25). Fundamental differences between fetal, infant, and adult gastrointestinal tracts raise questions on whether adult and fetal HIEs adequately model the infant gastrointestinal epithelium. To address the limited data on the morphological and functional characteristics of infant HIEs, we characterized jejunal HIEs from three infants and performed comparisons to HIEs from three adults. We also sought to determine if infant cultures recapitulate known features of infant intestinal epithelial biology and function such as lower barrier integrity and immune responses (28, 29). In this study, we demonstrate transcriptional, morphological, and functional differences between infant and adult HIEs supporting the use of infant HIEs to model infant-specific diseases.

## Results

### Transcriptional signatures of infant HIEs are distinct from adult HIEs

To determine the transcriptional profile of infant and adult HIEs, RNA sequencing (RNA-Seq) was performed on 5-day differentiated jejunal HIEs plated as monolayers on transwells. HIE lines were established using surgical samples from three infants (J1005, J1006, J1009) and three adults (J2, J3, J11). Patient demographic information and reasons for surgery are shown in Table 1. Principal component analysis (PCA) showed distinct clustering of the adult and infant HIEs (Fig. 1A). We identified a total of 1955 differentially expressed genes (DEGs, fold change >1.5, p-adj <0.05) in infant HIE lines compared to adult HIEs (Fig. 1B & 1C), including 796 genes that are up-regulated in infant HIEs compared to adult lines, and 1159 genes that are down-regulated. The most significantly upregulated genes in the infant HIEs include fatty acid binding protein 6 (*FABP6*) and solute carrier family 10 member 2 (*SLC10A2*) (Fig. 1C) that are involved in the binding and transport of bile acids. *FABP6* is also involved in the uptake, transport, and metabolism of fatty acids. Alpha-fetoprotein (*AFP*), which is known to be upregulated primarily in gestation was also upregulated in infant HIEs (30, 31). The most significantly downregulated genes were leukemia inhibitory factor receptor (*LIFR*), a component of cell-surface receptor complexes for multi-functional cytokines, and carcinoembryonic antigen-related cell adhesion molecule 7 (*CAECAM7*), a cell surface glycoprotein involved in cell-cell adhesion. Gene set enrichment analysis (GSEA) using the gene ontology biological processes compendium (GOBP) on all expressed genes showed 557 up- and 1933 down-regulated pathways. The top upregulated pathways included RNA processing, ribonucleoprotein complex biogenesis, and ribosome biogenesis while the top downregulated pathways included biological adhesion, cell-cell adhesion, and cell migration (Fig. 1D). Over-representation analysis (ORA) using only DEGs showed significant differences in regulation of cell differentiation and proliferation, lipid metabolism and immune response (Fig. 1E). Together, these data demonstrate striking differences in the transcriptome of HIEs derived from infants and adults.

**Table 1:** Demographic characteristics of HIEs from infants and adults. mo = months, wks = weeks, yo = years old, NA = not available.

| Line | Sex | Race | Ethnicity | Age at Surgery | Corrected Gestational Age at Surgery | Surgical Diagnosis (& Co-Morbidity) | Surgical procedure |
| --- | --- | --- | --- | --- | --- | --- | --- |
| J1005 | F | Caucasian | Non-Hispanic | 10 wks | 37 wks | Segmental ileal volvulus (Intrauterine growth restriction) | Jejunostomy takedown |
| J1006 | F | Caucasian | Non-Hispanic | 12 wks | 43 wks | Meconium peritonitis | Jejunostomy and mucus fistula takedown |
| J1009 | M | Caucasian | Hispanic | 22 wks | 54 wks | Jejunal atresia (Intrauterine growth restriction) | Jejunostomy and colostomy takedown |
| J2 | F | unknown | unknown | 52yo | NA | Morbid Obesity | Bariatric surgery |
| J3 | F | unknown | unknown | 46yo | NA | Morbid Obesity | Bariatric surgery |
| J11 | F | unknown | unknown | 52yo | NA | Bowel obstruction (Morbid Obesity) | Bowel resection |

**Figure 1:**
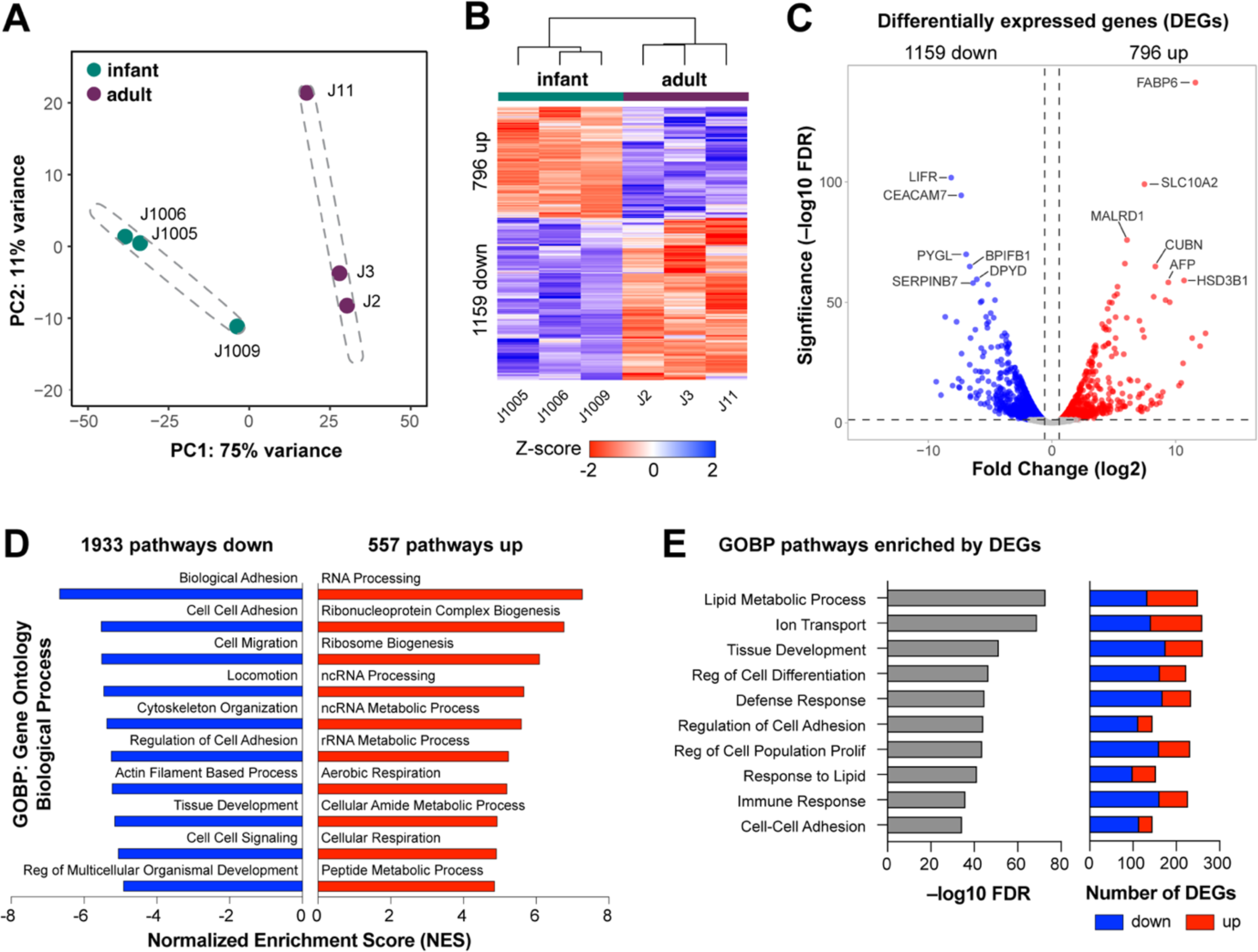
Infant HIEs have a different transcriptome than adults. A: Principal component analysis of infant (J1005, J1006, J1009) and adult (J2, J3 J11) HIEs, with the dashed ellipses indicating 95% confidence interval. B: Heatmap of genes differentially expressed in infant over adult HIEs (fold change exceeding 1.5, p adj.<0.05) C: Volcano plot of differential gene expression analysis results. Each gene is represented by a dot; grey dots represent no significant difference between infant and adult HIEs, the blue dots represent significantly down-regulated genes in infant HIEs, and red dots represent significantly up-regulated genes in infant HIEs. The top 12 genes by statistical significance are annotated. D: Gene set enrichment analysis (GSEA) results using the Gene Ontology (GO) Biological Processes compendium showing the top 10 up- and down-regulated pathways in infant over adult HIEs. E: Summary plot for over-representation analysis (ORA) showing 10 significantly enriched pathways selected for further validation. Number and direction of gene changes are shown in the plot on the right.

### Cell type composition of differentiated HIEs varies between infant and adult HIEs

Regulation of cell differentiation was significantly different between infant and adult HIEs (Fig. 1E). We first compared transcript expression for markers of absorptive (enterocytes) and secretory cells (goblet cells, enteroendocrine cells (EECs), Paneth cells and tuft cells) between infant and adult HIEs. Selected genes included alkaline phosphatase (*ALPI*), sucrase isomaltase (*SI*) and fatty acid binding protein 6 (*FABP6*) for enterocytes; trefoil factor 3 (*TFF3*), mucin 2 (*MUC2*) and chloride channel accessory 1 (*CLCA1*) for goblet cells; chromogranin A (*CHGA*), chromogranin B (*CHGB*) and somatostatin (*SST*) for enteroendocrine cells; Defensin Alpha 6 (*DEFA6*), regenerating family member 3 alpha (*REG3A*), and lysozyme (*LYZ*) for Paneth cells, and transient receptor potential cation channel subfamily M member 5 (*TRPM5*), advillin (*AVIL*), and POU class 2 homeobox 3 (*POU2F3*) for tuft cells (32). Except for *TFF3*, which was not significantly different, all markers for enterocytes, goblet cells and enteroendocrine cells were expressed at higher levels in infant HIEs compared to adult cultures (Fig. 2A). The transcript expression of Paneth and tuft cells markers had no consistent differences. Higher transcriptional expression of enterocyte, goblet and enteroendocrine markers corresponded to significantly higher abundance of cells expressing SI, MUC2, and ChgA (Fig. 2B and 2C) when HIE monolayers were stained for these markers. A striking observation from the confocal microscopy studies was the number of MUC2 positive cells in all three infant HIE lines. We stained sections from intestinal tissue samples used to generate infant HIEs to determine if high abundance of MUC2 expressing goblet cells was a feature of these tissues. Similar to the infant HIEs, we observed high MUC2 expression in infant tissue samples suggesting infant HIEs retain characteristics of the donor tissues (Sup. Fig. 1). Tissue sections from adult donors used to generate the adult HIE lines in this study were not available since they were some of the earliest cultures established at our organoid core. However, to determine if there are differences in goblet cell expression between adults and infant tissues, we stained three other adult tissue sections (J2001, J2002, J2003) and found overall higher MUC2 expression in infant tissues, thus validating differences observed between infant and adult HIEs (Sup. Fig. 1).

**Figure 2:**
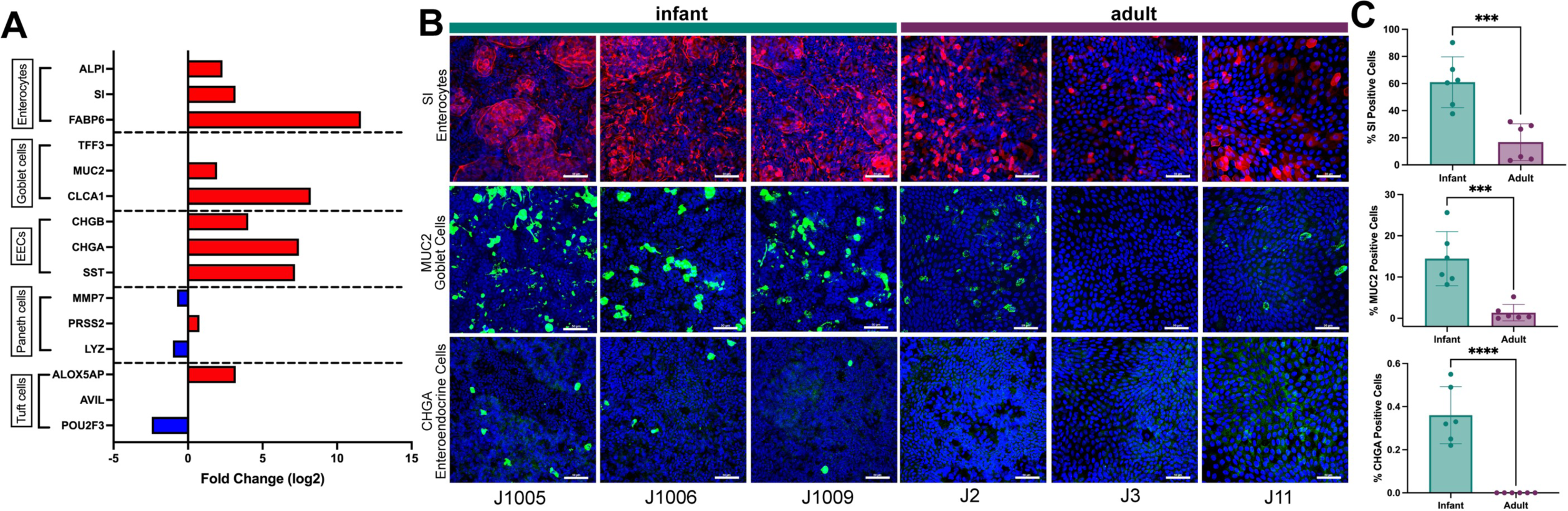
Cell type composition of varies between differentiated infant and adult HIE monolayers on transwells. A: RNAseq transcript expression for select markers of absorptive (enterocytes) and secretory cells (goblet cells, enteroendocrine cells, Paneth cells and tuft cells). Data represent mean values and are expressed as Log2 fold change of gene expression in infant over adult HIEs. Red bars = upregulated in infant HIEs; blue bars = downregulated in infant HIEs. B: Representative confocal images from four independent experiments, with each experiment including the three infant and three adult HIE lines. Top panel: enterocytes stained for expression of sucrase isomaltase (SI, red), middle panel: goblet cells (Muc2, green), and bottom panel: enteroendocrine cells (ChgA, green). Nuclei are stained with DAPI (blue), Scale bar = 50 μm. C: Quantification of cell type abundance from two independent experiments. Data represent mean ± SD with each experiment including the three infant and three adult HIE lines. The *p*-values were calculated by one-way ANOVA. The asterisks (***) and (****) represent p < 0.001, and p < 0.0001 respectively.

### Undifferentiated 3D infant HIEs are more proliferative than adult HIEs

Pathway analysis also showed differences in regulation of cell population proliferation between infant and adult HIEs (Fig. 1E). We first assessed 5-Ethynyl-2’deoxyuridine (EdU) incorporation on differentiated HIE monolayers plated on transwells by flow cytometry. We found a significantly higher percentage of proliferative cells in infant HIEs (9.9%) than adult HIEs (3.9%, p=0.02) (Sup. Fig. 2A & 2B). By virtue of being differentiated cultures, the numbers of proliferative cells are low; therefore, we also assessed EdU incorporation in undifferentiated infant and adult HIE monolayers plated on transwells. There were no significant differences in proliferation between infant and adult undifferentiated HIE monolayers (Sup. Fig. 2C & 2D). Next, to evaluate if HIE plating format influences the outcome of these assays, we assessed EdU incorporation in undifferentiated and differentiated 3Ds HIEs (Fig. 3 and Sup. Fig. 4, respectively). Immunofluorescence staining and flow cytometry quantification showed significantly higher proliferative cells in undifferentiated 3D infant HIEs (29.6%) than in adult HIEs (18.4%, p=0.03) (Fig. 3B). Donor specific differences in the percentage of proliferative cells were seen with the infant J1006 line being the most proliferative (35.7%) while the adult J2 HIE was the least proliferative (11.9%) (Fig. 3C). Compared to undifferentiated 3D HIEs, there were very few EdU positive cells in differentiated 3D HIEs (about 2%). There were no significant differences in percent EdU positive cells between differentiated infant and adult 3D HIEs (Sup. Fig. 3A & 3B). The contrast in results between HIE monolayers on transwells and 3D cultures suggest that plating format may be an important consideration when performing proliferation assays on HIEs.

**Figure 3:**
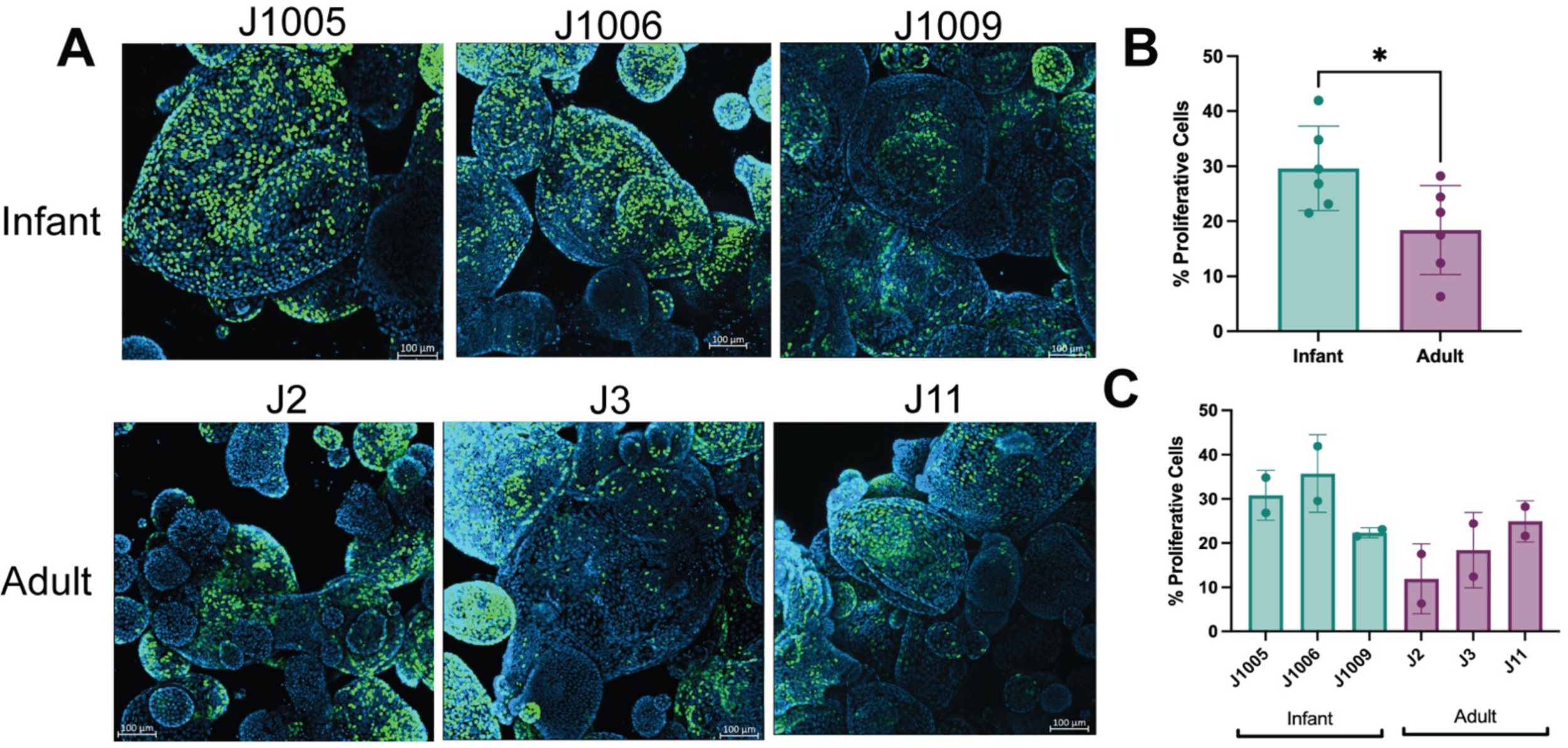
Undifferentiated infant 3D HIEs are more proliferative than adult HIEs. A: Representative confocal 3D reconstruction images after 24h 5-ethynyl-2′-deoxyuridine (EdU) incorporation in undifferentiated infant and adult HIEs. B: Percentage of EdU-positive cells quantified by flow cytometry using data combined from all infant or adult lines. C: Percentage of EdU-positive cells quantified by flow cytometry in all six lines. Data represents mean ± standard deviation (SD) from two independent experiments, with each experiment including the three infant and three adult HIE lines. The *p*-values were calculated by student’s t-test, and the asterisk (*) represents p < 0.05.

### Infant HIEs have shorter epithelial cell height and lower barrier integrity than adult HIEs

Pathway analysis showed differences in tissue development and downregulation of cell-cell adhesion in infant HIEs. These data suggest potential morphological differences as well as functional differences in barrier integrity. We first evaluated differences in the cellular morphology of HIE lines through measurement of single cell heights from H&E-stained five-day differentiated monolayers on transwells (Fig. 4B & 4C). Infant HIEs had significantly shorter cell heights than their adult counterparts (Fig. 4A). We also observed donor specific differences in cell height within both groups. Specifically, HIEs from J1009 were significantly shorter than the other two infant lines (Fig. 4E) and J3 was significantly shorter compared to the other adult lines (Fig. 4F). Similar to cell type composition studies, infant tissue sections were used to determine if HIEs reflect characteristics of donor tissues from which they are derived. Cell height measurements of the infant tissue (Fig. 4G) showed the same trend as the HIEs (Fig. 4E), with the cell heights from J1009 tissue section being significantly shorter than the other two infant tissues.

**Figure 4:**
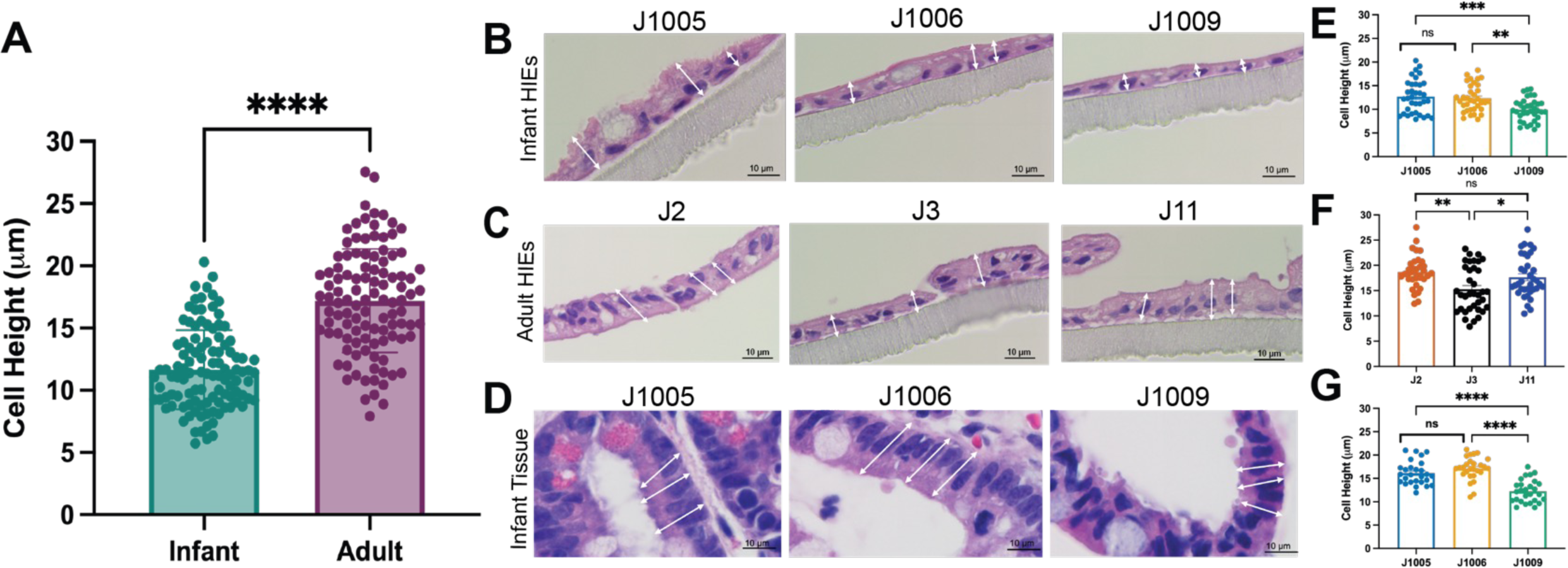
Infant HIEs have shorter epithelial cell height. A: Quantitation of cell heights measured in H&E-stained images of five-day differentiated infant and adult HIEs; *p*-values were calculated by student’s t-test. B-D: Three representative images were taken for each HIE line and tissue, and blinded images were provided to three study authors for cell height measurements. A minimum of three single cells per image were measured by each analyst, generating up to 35 data points per sample. Panel D shows paired donor tissue samples for each infant HIE line. E-G: Each dot indicates measurement of a single cell. Data represent mean ± SD with each experiment including the three infant and three adult HIE lines. The *p*-values were calculated by one-way ANOVA. The asterisks (**), (***), and (****) represent p < 0.01, p < 0.001, and p < 0.0001 respectively. Scale bar = 10 μm.

To further characterize differences in barrier function, we analyzed differences in tight junction markers in the RNA-Seq dataset. While there were no significant differences in the expression of tight junction protein (TJP) 1, 2 and 3 and occludin transcripts (Sup. table 1), several claudin genes were significantly downregulated in the infant lines compared to adult HIEs (Fig. 5A). However, claudin-2, a known marker of leaky epithelia (33), was significantly upregulated in the infant lines. To determine if transcriptional differences also result in changes in protein expression, we performed western blot analysis to quantify claudin-2 protein expression. Compared to adult HIEs, the infant HIEs had significantly higher expression of claudin-2 (p =0.005) (Fig. 5B) suggesting that infant HIEs are likely to be more leaky than adult lines. We therefore assessed barrier integrity of HIE monolayers on transwells by measuring TEER. The infant HIEs had significantly lower TEER compared to adult HIEs (p=0.0003) (Fig. 5C). TEER measurements varied between HIE lines although differences were not statistically significant (Fig. 5D).

**Figure 5:**
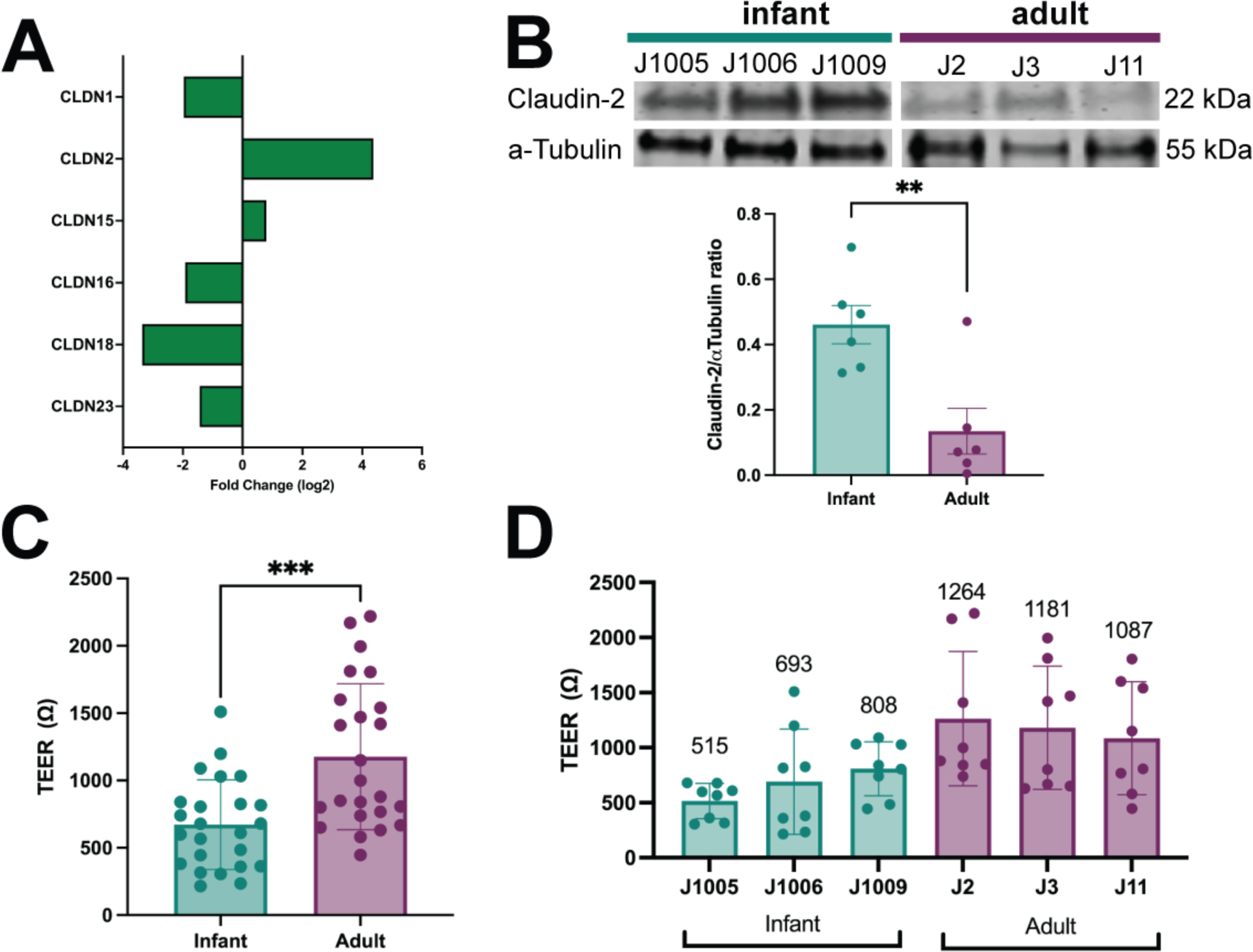
Infant HIEs have lower barrier integrity than adult HIEs. A: Log2 fold change of claudin genes expressed in infants over adults, as determined by RNA-Seq. B: Western blot of expressed CLDN2 in infant and adult HIEs (top) and densitometry-based quantification (bar plots) of CLDN2 normalized to α-Tubulin. C-D: Transepithelial electrical resistance values of five-day differentiated HIE transwell monolayers. Data represent mean ± SD of TEER measurements from four independent experiments, with two transwells/line. One measurement was taken for each transwell. The asterisks (**), (***) represent p < 0.01, p < 0.001 respectively.

Based on the lower TEER and higher claudin-2 expression observed in the infant lines, we next evaluated functional differences in permeability by measuring 4kDa fluorescein isothiocyanate (FITC)-dextran_translocation across the HIE monolayer. Cells were treated with ethylene glycol-bis(beta-aminoethyl ether)-N,N,N’,N’-tetra acetic acid (EGTA), a specific calcium chelator shown to have dramatic effects on paracellular permeability and TEER (34, 35). Treatment with EGTA caused a significant drop in TEER of both infant and adult HIEs (Fig. 6A). However, the infant HIEs (p=0.00004) were more sensitive to barrier disruption with EGTA treatment compared to the adult (p=0.001) HIEs (Fig. 6A). At baseline, there were no significant differences in FITC-dextran translocation across the infant (1.2μg/mL) HIE monolayers compared to adult HIEs (0.7μg/mL) (Fig. 6B). However, after EGTA treatment, we detected significantly higher FITC-dextran translocation in the infant HIEs (20μg/mL) compared to adult HIEs (6.5μg/mL, p = 0.04) (Fig. 6C) suggesting that infant HIEs may be more leaky following barrier disruptions. Donor specific differences in FITC-dextran translocation at baseline and with EGTA treatment are shown in (Sup. Fig. 4A & 4B). Tumor necrosis factor-α (TNFα), often used to simulate NEC *in vitro,* is also regarded as a modulator of epithelial permeability (36, 37). We tested the effect of TNFα on our HIE lines and found that basolateral administration of TNFα caused a significant drop in TEER in both infant and adult HIEs (Sup. Fig. 5A). However, contrasting the results with EGTA treatment, there was minimal FITC-dextran translocation in the infant and adult lines in response to TNFα (Sup. Fig. 5B).

**Figure 6:**
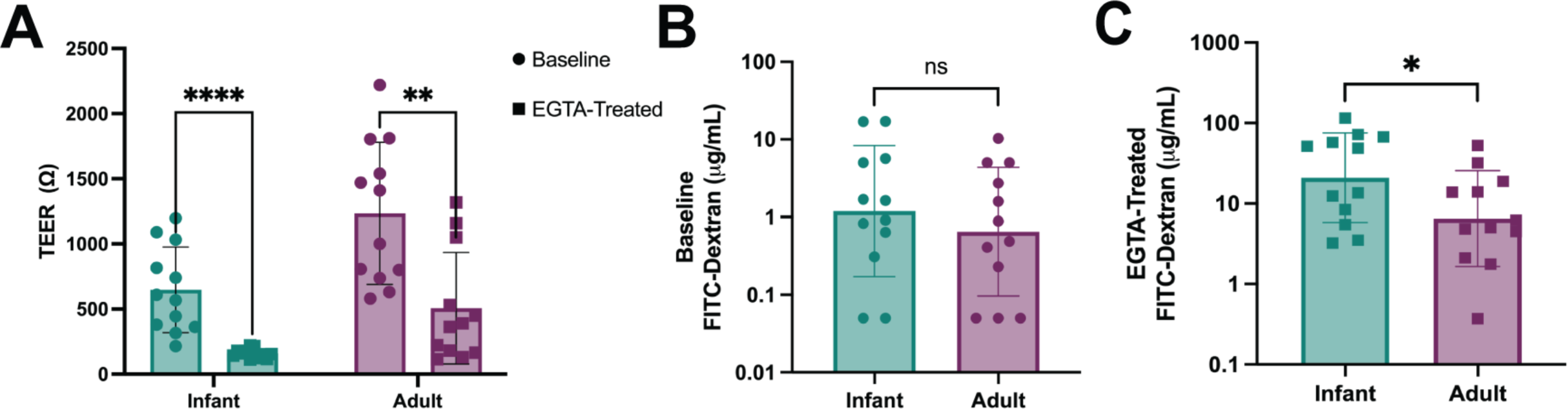
Infant HIEs have higher epithelial permeability in response to EGTA than adult HIEs. A: TEER values of HIEs at baseline and after EGTA treatment. B: Concentration of 4kDA FITC-Dextran at baseline and C: after EGTA treatment. Data represent mean ± SD from four independent experiments, with each experiment including the three infant and three adult HIE lines. The *p*-values were calculated by student’s t-test, and the asterisks (*), (**) and (****) represent p < 0.05, p < 0.01, and p < 0.0001 respectively.

### Infant HIEs express higher levels of lipid and lactose metabolizing genes

Pathway analysis of RNA-Seq data showed significant differences in lipid metabolic process and response to lipids (Fig. 1E). We evaluated the expression of genes involved in lipid absorption, trafficking, and metabolism by RNA-Seq (Fig.7A) and RT-qPCR (Fig. 7B-D). Infant HIEs had significantly higher expression of *MTTP* (codes for microsomal triglyceride transfer protein involved in triglyceride and cholesterol transport) and *APOB* (encodes Apolipoprotein B-48, a marker of intestinal chylomicrons) (Fig. 7B & 7C). RT-qPCR analysis also showed significantly higher expression of *LCT* (lactase) in infant HIEs (p <0.0001) (Fig. 7D). These data suggest that infant and adult HIEs may demonstrate differences in response to breast milk nutritional factors.

**Figure 7:**
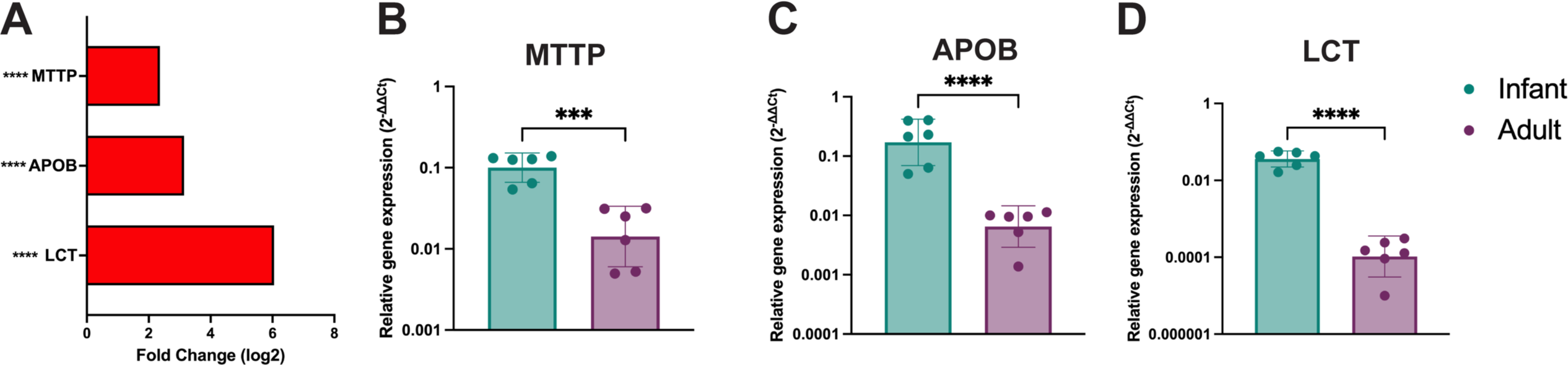
Lactase and lipid metabolism genes are significantly upregulated in infant HIEs. Expression profile of A: RNA-seq data for lactase and lipid metabolism genes B-D: RT-qPCR data for B: Microsomal triglyceride transfer protein (MTTP), C: Apolipoprotein B (APOB), D: Lactase (LCT) transcripts. Data represent mean ± SD from two independent experiments, with each experiment including the three infant and three adult HIE lines. The *p*-values were calculated by student’s t-test and the asterisks (***), and (****) represent p < 0.001 and p < 0.0001 respectively.

### Infant HIEs have lower interferon responses to an infant vaccine than adult HIEs

Pathway analysis of RNA-Seq data showed differences in immune response between infant and adult HIEs driven primarily by downregulated genes in infant cultures (Fig. 1E). To validate this observation, we evaluated innate epithelial immune responses to the monovalent type 1 oral poliovirus vaccine (mOPV1), a live, attenuated vaccine administered at birth to infants in low and lower-middle income countries (38). Both infant and adult HIEs supported similar replication of mOPV1, with about 3-log_10_-fold increase in the amount of infectious virus from 2 to 24 hours post infection (hpi) (Fig. 8A). Despite similar levels of replication, the expression of type III interferon; *IFNλ2* (interferon lambda) (Fig. 8B) and interferon-stimulated genes; *IFI44L*, (IFN-induced protein 44-like), and *IP-10*, (gamma-induced protein 10) (Fig. 8C & 8D) were significantly lower in the infected infant HIEs compared to adult lines. Donor specific differences in mOPV replication and the subsequent immune responses are shown in Sup. Fig. 6A-6D.

**Figure 8:**
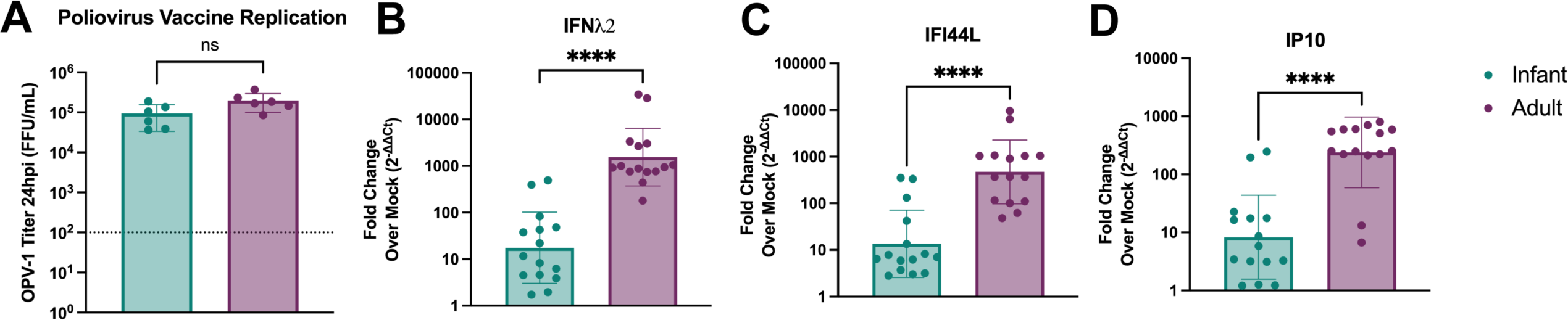
Infant HIEs have lower epithelial immune responses to oral vaccines than adult HIEs. A: Monovalent type I poliovirus vaccine (mOPV1) titer in HIEs at 24hpi. The line at 100 FFU/mL represents OPV-1 titer at 2hpi (baseline). Quantification of (B) IFNλ2, (C) IFI44L, and (D) IP10 transcripts. Data represent mean ± SD from two independent experiments, with each experiment including the three infant and three adult HIE lines. The *p*-values were calculated by student’s t-test and the asterisk (****) represents p < 0.0001.

### Infant and adult HIEs exhibit line-specific differences in response to an enteric virus

To determine if the differences observed in innate immune responses to mOPV1 are also seen for other enteric viruses, we infected the HIEs with human norovirus (HuNoV), a leading cause of acute gastroenteritis across all age groups. We first performed 50% tissue culture infectious dose (TCID_50_) assays on each HIE line with a globally dominant GII.4 HuNoV strain. Overall, HuNoV TCID_50_ values were lower for infant HIEs (average 7974 genome equivalents) compared to adult HIEs (average 12297 genome equivalents). However, the differences were not statistically significant (Sup. Fig. 7A) primarily because the TCID_50_ value for J1009 was similar to that of the adult HIEs (Sup. Fig. 7A). We then used a standard dose of virus (100 TCID_50_) to compare innate immune responses between different HIE lines. Infection with 100 TCID_50_ of HuNoV resulted in lower expression of *IFNλ2* and *IP10* and similar expression of *IFI44L* (Sup. Figs. 7B-D) in the infant HIEs although these differences were not statistically significant. This could also be attributed to HIE line-specific differences in innate immune responses to HuNoV infection (Sup. Fig. 8B-D). Of note, despite infection with uniform TCID_50_ of HuNoV, two of three infant HIE lines showed lower expression of innate immune response genes compared to the adult lines.

## Discussion

The intestinal epithelium of infants undergoes rapid changes in response to growing nutritional requirements and changes in the intestinal milieu. There are many differences in morphology and gastrointestinal function from birth to six months of age when compared to the adult gut (39, 40). The lack of age-appropriate and biologically relevant model systems has limited advances in mechanistic studies on the human infant intestinal development, nutrition, microbiome, and disease pathogenesis. Our in-depth transcriptional, morphological, and functional comparisons of HIEs from infants and adults suggest that infant HIEs recapitulate several unique *in vivo* characteristics of the infant epithelium and are important new tools for basic and translational studies of gastrointestinal development and diseases in this population.

The transcriptional and morphological analyses show that infant HIEs model the cell composition and morphological characteristics of the immature infant intestinal epithelium. Current literature mapping the cellular composition of human embryonic, fetal, and pediatric intestinal tissue report more enterocytes in the pediatric gut (41). While RNA-Seq analyses of fetal enteroids have been carried out in some studies, direct comparisons to our present work were limited because of differences in intestinal segment of derivation or media used for treatment which are both major drivers of variance in transcriptional responses (24, 42, 43). However, like fetal HIEs, the infant HIEs in our study display markers of the immature intestinal epithelium such as higher paracellular permeability and host-defense functions compared to adult HIEs (24). Single cell RNA-Seq and proteomic analysis of swine neonatal ileal epithelium, a widely used model of GI development because of difficulties in studying healthy human infant intestinal tissues, showed that the expression of goblet cells, EECs and tufts cells were highest at birth while the expression of enterocytes increased dramatically from birth, peaking at day 7 of life (44). Together, these data suggest high expression of both absorptive and secretory cells in the infant gut. The differentiated infant HIEs used in our studies showed high expression of specific enterocyte, goblet, and EEC markers similar to observations from pediatric tissues and the neonatal swine model. While we did not observe transcriptional differences in the expression of Paneth and Tuft cell markers, it is possible that there are differences in abundance of these cell types that could be analyzed using immunofluorescence staining or Western blot.

Other morphological characteristics of infant HIEs such as shorter epithelial cell height and higher proliferation also reflect features of a developing infant intestinal epithelium. Shorter epithelial cell height is described as a marker of intestinal immaturity and has been reported in duodenal biopsies of infants (45) and more recently, duodenal pediatric HIEs (25). Increased proliferation within the intestinal crypt contributes to crypt hyperplasia and is a potential indicator of an immature intestinal epithelium (46). Early studies of duodenal biopsies from infants and adults reported that crypt hyperplasia is specific to infancy and is a mechanism of postnatal growth of the human small intestine (47). Crypt fission, another mechanism of intestinal growth related to increased proliferation, was also reported to be highest at infancy and declined after 3–4 years (48). We observed significant differences in HIE proliferation in the 3D format. Surprisingly, undifferentiated infant and adult HIE monolayers on transwells showed no differences in proliferation. We recently reported that common differences in culture conditions such as substrate (collagen vs. Matrigel) and format (3D, transwell, and monolayer) are the largest drivers of transcriptional variance in HIEs (43). Our data extends these findings and suggest that plating format may impact some assays such as measurements of proliferation. On the other hand, HIE format did not influence assays such as immunofluorescence staining of cell types as the striking differences between infant and adult HIE monolayers in expression of MUC2 and CHGA were also observed in 3D HIEs (Sup. Fig. 9).

Our functional studies highlight the value of these physiologically- and developmentally appropriate models when studying infant-specific disease conditions. A key feature associated with many neonatal gastrointestinal diseases is barrier dysfunction. Breakdown of barrier function also plays a key role in the pathogenesis of infectious gastroenteritis and inflammatory bowel diseases and may predispose vulnerable population to sepsis (49, 50). The immature gut barrier has been well described as a predisposing factor in the pathogenesis of NEC in preterm infants (51–53). Modeling the epithelial barrier of an infant gut is therefore critical for studies of infant-specific diseases and testing interventions. Increased *TLR4* signaling has been shown to initiate a strong innate immune response that leads to epithelial barrier disruption and development of NEC (54, 55). Infant HIEs show significantly higher expression of *TLR4* (Sup. Fig. 10) and are sensitive to barrier disruptions. EGTA and TNFα resulted in a drop in TEER; however, while barrier disruption with EGTA led to significant increases in permeability of infant HIEs to FITC-dextran, this result was not observed with TNFα treatment. The differences in FITC-dextran permeability results suggest changes in ion transport as measured by TEER do not necessarily translate to similar changes in translocation of macromolecules (56).

Promising therapies to decrease the risk of NEC involves improving barrier function in the infant gut using growth factors (57, 58), breast milk and its bioactives (59), and probiotics (60–62). Indeed, a previous study using duodenal HIEs from a two-year-old and five-year-old child showed that colostrum enhanced epithelial barrier function and breast milk increased the production of alpha defensin 5 (25). Addition of breast milk was also shown to increase cell height and promote growth of these pediatric duodenal HIEs (25). While we did not directly evaluate the effects of breast milk on infant and adult HIEs, we examined genes involved in the metabolism of lactose and fats, the two most abundant solid components in human breast milk (50, 63). Lactase, the enzyme responsible for the breakdown of lactose to glucose and galactose, is expressed on the brush border of the small intestine. Lactase levels are reported to be higher in neonates than adults (63, 64). Similar observations were made in our infant HIEs when compared to adult HIEs. After carbohydrates, lipids are the second most abundant macronutrient and provide a major portion of the total energy needed by the infant (63, 64) and we found higher expression of genes such as *MTTP* and *APOB* involved in lipid absorption, trafficking, and metabolism. Majority of milk lipids are triacylglycerols and their properties depend on the incorporated fatty acids (65). Human milk also contains significant amounts of cholesterol (66). The most significantly upregulated gene in infant HIEs was *FABP6*, which is involved in the uptake, transport, and metabolism of fatty acids. Additionally, several lipid metabolism genes were upregulated in the infant HIEs, suggesting that these cultures could be used for studying nutritional factors in breast milk.

The innate immune system provides an early first line of defense against invading pathogens. While immune cells develop and mature during fetal life, the function of all components of innate immunity are weaker in infancy compared with later life (67–71). For example, neutrophils, monocytes, and macrophages have weaker antimicrobial functions and innate signaling pathways compared to adults (72–75). Differences in immune response associated genes has previously been reported in comparison of preterm and adult ileal HIEs (26). We show that infant HIEs exhibited lower epithelial innate immune responses on infection with a live, attenuated mOPV1 vaccine that is administered as a birth dose. Prematurity could therefore explain the lower immune responses in the infant lines as the preterm infant innate immune system is characterized by attenuated pro-inflammatory and antiviral cytokine responses (70, 76). Surprisingly, despite the lower immune response in the infant HIEs, OPV replicated similarly in both adult and infant lines. We surmise that this may be attributed to high multiplicity of infection (MOI) used to infect the HIEs or the higher numbers of susceptible enterocytes in infant cultures. Compared to our results with mOPV1, there was more variability in response to HuNoV infection that causes infections in all age groups (77). Lower TCID_50_ values and lower innate immune responses were observed with two of three infant HIEs (J1005 and J1006), suggesting that those infant lines may be more susceptible to HuNoV infection compared to J1009 and the adult HIEs. Why the striking differences in mOPV1 innate immune responses were not uniformly observed with HuNoV infection remain to be elucidated. Lower innate responses may continue beyond the neonatal period as demonstrated by a study that showed pediatric duodenal HIEs from two- and five-year old children did not produce TGF-b1, IFN-g, and IL-6 which have been detected in adult HIEs (25, 78). Our results suggest that infant lines accurately model the innate immune status of a developing gut and that it may be more relevant to use age-appropriate models when studying enteric infections and oral vaccines. These data also suggest innate epithelial development HIEs from infants and children of different age groups can be used to model innate epithelial development, especially as co-culture models with immune cells continue to be developed (78, 79).

In addition to recapitulating many unique characteristics of the infant intestinal epithelium, HIEs also reflect specific characteristics of the hosts from which they are established. This is evidenced by the MUC2 staining and intestinal tissue height measurements for infant HIEs. However, we observed donor-specific differences between the different infant HIE lines in multiple experiments. For example, the J1009 HIE line had a lower percentage of proliferative cells, fewer numbers of EECs, lower cell height and higher TEER than the other two infant lines (Figs. 2B, 3C, 4B, 4D, 4E, 4G). This may be due to the age of the host, since J1009 HIE was established from a 5-month-old infant (corrected gestational age 52 weeks) and is therefore an older donor than the other two lines which were collected at 2-3 months of age. There were also ethnicity and biological sex differences between J1009 and other lines. Whether and how any of these factors contribute to differences between donors is difficult to elucidate given the small sample size of our study. The differences in barrier integrity and intestinal permeability may be attributed to the feeding status of the donors; however, since all samples were deidentified at the time of establishing HIEs, feeding data are currently not available for analysis. Expanding the sample size of HIEs lines used for comparison and expanding on metadata collection in future studies will help address if any of these factors contribute to differences observed. None-the-less, these studies demonstrate how HIEs model the heterogeneity of the human population and highlight study design challenges when considering sample size calculations.

Studies of intestinal pathogens and disorders of the human gastrointestinal tract have traditionally used transformed or immortalized *in vitro* culture systems that have limited physiological relevance to the human intestinal epithelium. In new legislation (80), the U.S. Food and Drug Administration no longer requires new medicines to be tested in animals and encourages the use of computer modeling, organ-on-chips and non-animal testing methods. As HIEs continue to become the standard biological model to address questions about host-pathogen interactions and for drug discovery, there’s a need to expand our bank of cultures to include donors from different ages, developmental stages, disease conditions and intestinal segments. There is also a need to increase complexity of these cultures by incorporating components of the immune system, enteric nervous system, and microbiome. A first step towards these goals is in-depth characterization of HIEs. Our studies provide an example of such characterization and further show that infant HIEs are age-relevant models that should be used in studies of infant intestinal physiology, mechanisms of infant-specific diseases, and as a platform for drug discovery for infants.

## Methods

### Human intestinal enteroids

Jejunal tissue samples were collected from infants undergoing gastrointestinal surgery following approval by the University of Texas Health Science Center at Houston Institutional Review Board (IRB) and parental consent. Infant tissue samples were transferred to Baylor College of Medicine under a material transfer agreement for establishment of HIEs. Adult jejunal tissue was obtained under an IRB-approved protocol at Baylor College of Medicine from patients undergoing bariatric surgery (16). HIE cultures were generated at the Texas Medical Center Digestive Diseases Center Gastrointestinal Experimental Model Systems Core from intestinal crypts isolated from the surgical tissues of infant and adult patients as previously described (81, 82). HIE monolayers used for this manuscript were all plated at a density of 2.5X10^5^ cells per transwell. Demographic details including age at sample collection and reasons for surgery are provided in Table 1.

### RNA Sequencing

We performed RNA-Seq analysis on five day-differentiated jejunal HIEs plated as monolayers on transwells. Total cellular RNAs were extracted using the RNeasy Mini Kit (Qiagen). RNA purity and concentration were measured using Nanophotometer Pearl (IMPLEN). Raw sequence reads were checked for quality using the FASTQC package ver. 0.11.9; Illumina adapters and low-quality basepairs were trimmed using TrimGalore ver. 0.6.5 with default settings. Trimmed reads were aligned to human genome build GRCh38.98 using HiSAT2 ver 2.2.1 (83) and a count matrix was generated from the aligned reads using featureCounts (84). Differential gene expression analysis was performed for the protein coding genes using the EdgeR ver. 3.32.1 R package (84). Differential gene expression significance was achieved for false discovery rate (FDR)-adjusted p-value<0.05 and fold-change exceeding 1.5x. Sequencing data and raw count matrix files were deposited in the Gene Expression Omnibus GSE227205. Enriched pathways for each comparison of interest were determined using Gene Set Enrichment Analysis (GSEA) v 3.333 (85) using the Gene Ontology pathway compendium compiled by the MsigDB database (86). The GSEA analysis was performed on rank files comprised of gene symbols and the corresponding log2 fold changes for all the expressed genes; enrichment was considered significant for adjusted *p*-value FDR<0.25. Functional analyses on the differentially expressed genes were further performed using Over Representation Analysis (ORA) (87) to determine whether known biological functions or processes are over-represented. We used the hypergeometric distribution as implemented by MSigDB, with significance achieved for adjusted *p*-value FDR<0.05.

### RNA Extraction and RT-qPCR

Total RNA was extracted using Direct-zol™ RNA MiniPrep Kit (Zymo Research) according to the manufacturer’s protocol. PCR reactions were performed using the qScript XLT One-Step RT-qPCR ToughMix reagent with 6-carboxy-X-rhodamine (ROX) (Quanta Biosciences) in the StepOnePlus real-time PCR system. Fold changes in mRNA expression were determined using the delta-delta-Ct method relative control samples after normalization to the housekeeping gene glyceraldehyde3-phosphate dehydrogenase (GAPDH). The following TaqMan primer-probe mixes (Thermo Fisher Scientific) were used: Hs99999905_m1 (glyceraldehyde3-phosphate dehydrogenase, GAPDH), Hs00820125_g1 (IFNλ2, IFNL2), Hs00915292_m1 (IFN-induced protein 44-like, IFI44L), Hs00171042_m1 (Interferon gamma-induced protein 10, *IP-10*), Hs00158722_m1 (Lactase, LCT), Hs0155363_m1 (Microsomal triglyceride transfer protein*, MTTP),* Hs00181142_m1 (Apolipoprotein B, *APOB)*

### Cell Height Measurements

Tissue sections and five day differentiated HIEs monolayers on transwells were fixed with 4% paraformaldehyde overnight at 4°C, washed with 1X PBS and immersed in 70% ethanol. Paraffin embedding, sectioning, and hematoxylin and eosin (H&E) staining were performed at the Human Tissue Acquisition and Pathology Core in Baylor College of Medicine. Three representative images were taken for each HIE line and tissue, and blinded images were provided to three study authors for cell height measurements. Cell heights were measured from the base of the cell to the top of the cilia; cells that were stacked in multiple layers and those that contained inclusions, vacuoles, or holes were excluded. Each author measured a minimum of three single cells per image using the Fiji ImageJ software, generating up to 35 data points per sample.

### TEER Measurement and FITC-dextran Permeability Assay

HIEs were plated as monolayers on transwells and differentiated for five days. TEER was measured before EGTA treatment to obtain baseline values. Cells were treated with 5mM EGTA from a 0.2M stock solution in distilled water that was pH adjusted to 7.4 with 1M Tris-HCl. EGTA was added to the apical side of the transwells and incubated at 37°C for 2h. After 2h, EGTA was removed and FITC-labeled 4 kDa dextran was added apically. For the permeability assay with TNF-α, HIEs were plated as monolayers on transwells and differentiated for four days. TEER was measured before TNF-α treatment to obtain baseline values. 0.25μg/mL of TNF-α was prepared in differentiation media and added to the basolateral side of the HIE transwells for a 24h treatment period. After 24h, TEER was measured and FITC-dextran added to the apical side of the HIE transwells. For preparation of FITC-dextran solution, 5mg of 4 kDa FITC-dextran was dissolved in 1mL of HIE differentiation media and filtered with a 0.2mm PVDF filter membrane. The 5mg/mL FITC-dextran was further diluted 1:100 in differentiation media, added to the transwells apically, and incubated for 2h. After 2h, media from the apical and basal side of each transwells was collected and assayed in duplicates using a fluorescent plate reader at an excitation/emission wavelength of 485/538nm. Average fluorescence intensity values were compared to a standard curve ranging from 0 to 5 mg/mL 4-kDa FITC-dextran prepared in differentiation media. FITC-dextran concentration (μg/mL) in the basolateral media was extrapolated from the standard curve using GraphPad Prism. TEER was measured again at the end of FITC-dextran treatment. A control (no EGTA or TNF-α treatment) was used in all experiments.

### Immunofluorescence Staining and Confocal Imaging

HIE monolayers on transwells were fixed with 4% paraformaldehyde in PBS pH 7.4, for 30 min at room temperature. Permeabilization was performed using 0.5% Triton X-100 in PBS, for 20 min at room temperature and with a blocking step consisting of 2% BSA in 0.1% Triton X-100 in PBS, for 30 min at room temperature. Primary antibodies targeting CHGA (1:1000 dilution; Immunostar, 20085), MUC2 (1:50 dilution; Santa Cruz, sc-515032), or SI (1:5 dilution; Developmental Studies Hybridoma Bank, HBB2-219-20) were added to the membranes and incubated overnight at 4°C. The next day, membranes were washed with PBS three times for 5_min each and species-specific secondary antibodies were added at a dilution of 1:1000 and incubated overnight at 4°C. Nuclei were detected by incubating in DAPI for 15 min at room temperature. The membranes were washed with PBS three times for 5_min each and mounted on glass slides using ProLong Gold antifade mounting medium. Paraffin embedded infant tissue sections were deparaffinized in Histoclear and rehydrated with successive ethanol baths. Antigen retrieval was performed with 10mM sodium citrate pH 6.0 for 1 minute in a pressure cooker and blocked with 1x PowerBlock (Biogenex). Slides were stained with MUC2 (1:100 dilution; Abcam, ab272692) at 4°C for 24h, washed in PBST (0.05% Tween20/PBS), and incubated with secondary antibody at room temperature for 1h. Slides were mounted with VECTASHIELD (Vector Labs). Immunofluorescence images were acquired using a Nikon A1Rs confocal laser scanning microscope using sequential image acquisition with laser power set between 5 and 10%. The pinhole was set at 1.0 for the 488nm or 561nm laser and the gain for any given image ranged from 85 to 130 (maximum possible gain, 255). All observations were carried out at the Integrated Microscopy Core in Baylor College of Medicine. Image analysis was performed on grey scale images using a custom-built pipeline in Cell Profiler 4.2.1. In brief, the nuclei were detected after applying a Gaussian filter for smoothing using Otsu thresholding. The nuclear mask was then expanded by 10 pixels and fluorescence intensity was extracted from the antibody channel. Single cell data was then evaluated and a threshold for positive cells was established via visual evaluation of a subset of images/antibody.

### EdU Proliferation Assay

EdU is a thymidine analogue that is incorporated into DNA as it is being replicated and is actively taken up into proliferating cells during the S-phase of the cell cycle. EdU labeling was carried out using the Click-iT EdU Alexa Fluor 488 Imaging Kit (Life Technologies). Undifferentiated and differentiated 3D HIEs or HIE monolayers on transwells were treated with 10mM EdU for 24h at 37°C to label proliferating cells. For immunofluorescence staining, 3D HIEs and HIE monolayers fixed with 4% paraformaldehyde for 20_min, washed with 3% bovine serum albumin and permeabilized using 0.5% Triton X-100. EdU reaction was carried out according to manufacturer’s protocol. The cells were visualized using LSM 980 confocal microscope (ZEISS) and images were taken using a 10X objective. For flow cytometry quantitation, 3D HIEs were dissociated into single cells by washing the cells out of the Matrigel plug and incubating the cells in cold Cell Recovery Solution (Corning) on ice for 10 minutes. Cells were spun at 400g for 3 minutes, dissociated using Accutase (Sigma) and incubated at 37°C for 30 minutes. Cells were gently pipetted every 10 minutes. After 30 minutes, the cells were pelleted at 400g for 3 min and resuspended in 1mL of complete medium without growth factors (CMGF-). This mixture was pipetted up and down 30 times rapidly to dissociate the cells. Finally, the suspension of cells was passed through 40μm and 70μm filters (VWR) and washed with 1% BSA/PBS. For flow cytometry quantitation, 0.05% trypsin was added to dissociate HIE monolayers from the transwell membranes and incubated for 5min at 37°C. Dissociated EdU labeled 3D HIEs and monolayers were fixed, permeabilized, and stained using the Click-iT™ EdU Alexa Fluor™ 488 Flow Cytometry Assay Kit (Thermo Fisher Scientific). Alexa Fluor 488-positive cells were quantified by using an LSRII flow cytometer (BD Biosciences). Doublet discrimination was used to gate on the single-cell population. From this gate, 10,000 events were analyzed (Sup. Fig. 11). Flow cytometry analysis was carried out at Cytometry and Cell Sorting Core in Baylor College of Medicine.

### Western Blot

Five-day differentiated 3D HIEs were solubilized in radioimmunoprecipitation assay buffer (Thermo Fisher Scientific) containing protease inhibitors (complete, mini, EDTA-free protease inhibitor cocktail tablets; Sigma Aldrich, 11836170001) and benzonase per the manufacturer’s instructions. Cells were lysed on ice for 20 mins and pelleted by centrifugation at 10,000 rpm (13,800g) in a Beckman Coulter J2-HS centrifuge. The cell lysates were frozen at -20°C until used. Cell lysates containing 25μg of protein were prepared with Laemmli sample buffer containing β-mercaptoethanol for western blot analysis. Samples were heated for 10 mins at 70°C and loaded onto 4-20% polyacrylamide gradient gels (Bio-Rad) for electrophoretic separation in Tris-glycine-SDS buffer (Bio-Rad). Following electrophoresis, the proteins were transferred to nitrocellulose membranes using the iBlot 7-min blotting system (Thermo Fisher Scientific). Membranes were blocked for 1 h at room temperature in 1 x casein blocking buffer (Sigma) and immunoblotted using primary antibody to Claudin-2 (1:500 dilution; Thermo Fisher Scientific, 32-5600) and α-Tubulin (1:1500 dilution; Sigma-Aldrich, T6-199) followed by fluorescent secondary anti-mouse antibodies (1:10,000 dilution, LI-COR) in 1 x casein blocking buffer. Blots were washed three times with PBS– 0.5% Tween 20 and once with PBS prior to visualization and quantification using an Odyssey infrared imaging system (LI-COR). Band densitometry for Claudin-2 and αTubulin was determined in Image Studio software. For each HIE line, the Claudin-2 signal was normalized to respective αTubulin signal and plotted as the ratio of Claudin-2 to αTubulin signal.

### Poliovirus vaccine infection and determination of viral replication

Monovalent type 1 attenuated oral poliovirus vaccine (mOPV1) virus stocks were propagated in HeLa cells as previously described (87). Virus stocks were used to infect 5-day differentiated HIE monolayers on transwells at an MOI of 2. Briefly, mOPV1 was added to HIEs for 2h, washed 2 times with CMGF(-) medium to remove the inoculum, and incubated in differentiation medium for 24h. Supernatants were collected from infected HIEs at 2hpi and 24hpi. MA104 cells plated in 96-well plates to confluency were used to quantify the titer of the infectious mOPV1 from HIEs as follows. After 1h adsorption, the inoculum was removed, and cells were washed twice with Dulbecco’s Modified Eagle Medium. Infection was allowed to continue at 37°C for 7h. Infected MA104 cells were fixed with cold 100% methanol for 20 min at room temperature. The cells were washed with 1X PBS and incubated with anti-poliovirus 1 antibody, clone 583-G8-G2-A4 (1: 600 dilution, Millipore Sigma) at 37°C for 2h. The cells were washed 3 times with 1X PBS and incubated with goat anti-mouse IgG (H&L)-Alexa Fluor 488 (1:1000 dilution, Thermo Fisher Scientific) at 37°C for 2h. Infected cells were counted using an epifluorescence microscope.

### Human norovirus infection

TCID_50_ values were determined for GII.4 Sydney[P16] (isolate BCM 16-16) as described previously (88, 89). To assess innate immune response to norovirus, 100 TCID_50_ of the virus in Intesticult media supplemented with 500 μM sodium glycochenodeoxycholate (GCDCA; Sigma, G0759) was added to 5-day differentiated HIE monolayers in triplicate on a 96-well plate. Each well had 4.26×104 genomic equivalents (GE)/well of virus. After incubating at 37°C for one hour, the inoculum was removed and the HIE monolayers were washed twice using CMGF-to remove unbound virus. 100ul intesticult media supplemented with GCDCA was added to the monolayers and incubated at 37°C for 23 hours. Total RNA was extracted using the Kingfisher Flex machine and MagMAX-96 viral RNA isolation kit as described previously (89).

### Statistical Analysis

Statistical analyses were performed using Prism software v9 (GraphPad). Statistical significance was determined using the student’s t-test or one-way ANOVA. Differences were considered statistically significant at p-value ≤ 0.05. All authors had access to the study data and reviewed and approved the final manuscript.

## Abbreviations

(*AFP*): Alpha-fetoprotein
(ALPI): alkaline phosphatase
(APOB): Apolipoprotein B-48
(*CAECAM7*): carcinoembryonic antigen-related cell adhesion molecule 7
(*CHGA*): chromogranin A
(CHGB): chromogranin B
(CLCA1): chloride channel accessory 1
(+/−): complete media with/without growth factors CMGF
(DEFA6): defensin alpha 6
(DEG): differentially expressed gene
(EdU): 5-Ethynyl-2’deoxyuridine
(EECs): enteroendocrine cells
(EGTA): ethylene glycol-bis(beta-aminoethyl ether)-N,N,N’,N’-tetra acetic acid
(*FABP6*): fatty acid binding protein 6
(FDR): false discovery rate
(FITC): fluorescein isothiocyanate
(GSEA): gene set enrichment analysis
(GOBP): gene ontology biological processes compendium
(HIEs): human intestinal enteroids
(IFNλ): interferon lambda
(IFI44L): IFN-induced protein 44-like
(IP-10): gamma-induced protein 10
(LCT): lactase
(*LIFR*): leukemia inhibitory factor receptor
(LYZ): lysozyme
(mOPV1): monovalent type 1 oral poliovirus vaccine
(ORA): over-representation analysis
(MOI): multiplicity of infection
(MTTP): microsomal triglyceride transfer protein
(*MUC2*): mucin 2
(NEC): necrotizing enterocolitis
(PCA): principal component analysis
(POU2F3): POU class 2 homeobox 3
(REG3A): regenerating family member 3 alpha
(*SLC10A2*): solute carrier family 10 member 2
(*SI*): sucrase isomaltase
(SST): somatostatin
(TLR): toll-like receptor
(TEER): transepithelial electrical resistance
(TFP): tight junction protein
(TFF3): trefoil factor 3

## Disclosures

The authors disclose no conflicts of interests.

### Data Transparency

RNA-Sequencing data files are deposited in Gene Expression Omnibus GSE227205. Other data that support the findings of this study are available from the corresponding author on request.

## Acknowledgements

This work was supported by funds from the National Institutes of Health (NIH) grants R21-A1132985 (SR), PhRMA Foundation Research Starter Grant in Translational Medicine (SR), PO1-AI057788 (MKE), U19 AI144297 (MKE) and U19 AI116497 (MKE) and the Texas Medical Center Digestive Diseases Center P30 DK056338. SLG and CC were partially supported by The Cancer Prevention Institute of Texas (CPRIT) RP170005, RP210227, RP200504, NIH P30 shared resource grant CA125123, NIEHS grants P30 ES030285 and P42 ES027725, and NIMHD P50MD015496. ALS was supported by NIH/NIDDK 1K08DK131326-01. This project was also supported by the Office of The Director, NIH under Award Number S10OD028480, the Advanced Technology Core Laboratories at Baylor College of Medicine, including the Integrated Microscopy Core at Baylor College of Medicine and the Center for Advanced Microscopy and Image Informatics (CAMII) with funding from NIH (DK56338, CA125123, ES030285, S10OD030414), and CPRIT (RP150578, RP170719), and the Cytometry and Cell Sorting Core with funding from the CPRIT Core Facility Support Award (CPRIT-RP180672) and the NIH (CA125123 and RR024574) and the assistance of Joel M. Sederstrom. We thank Dr. Richard Lloyd for providing the monovalent poliovirus vaccine.

