## Supplemental Figure 1 for "Infant and Adult Human Intestinal Enteroids are Morphologically and Functionally Distinct"

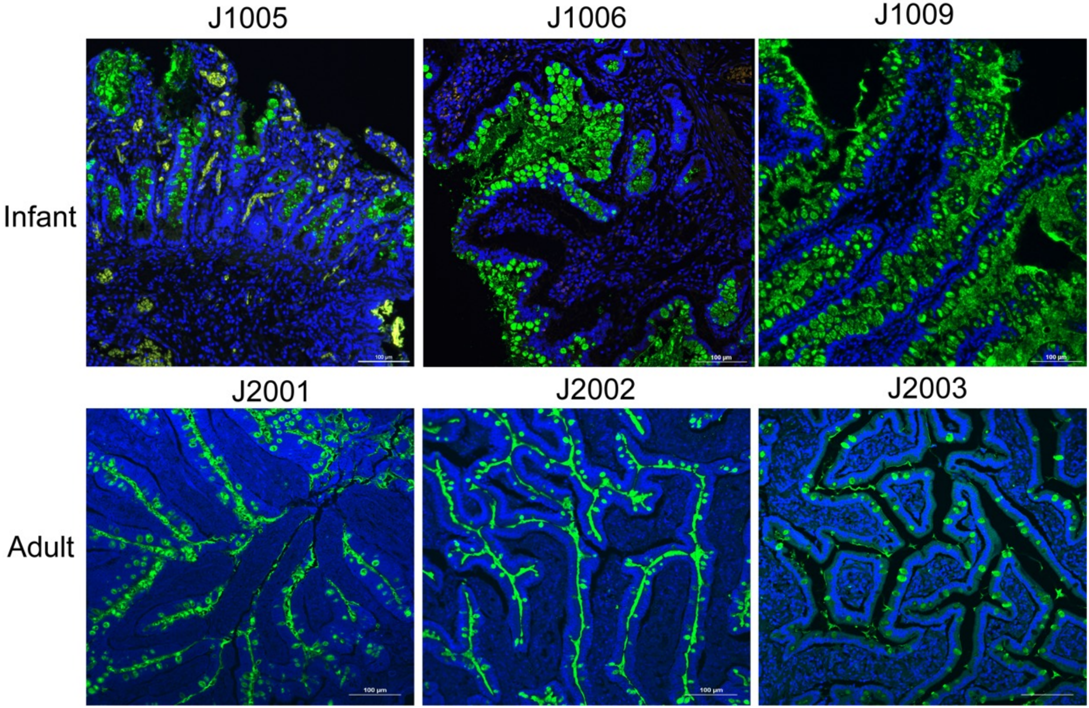


**Supplemental Figure 1: MUC2 expression is higher in infant intestinal tissues than adult tissues**

Representative confocal images of goblet cells (Muc2, green) in infant and adult tissues from two independent experiments. Nuclei were stained with DAPI (blue). Scale bar = 100 µm.
