## Supplemental Figure 2 for "Infant and Adult Human Intestinal Enteroids are Morphologically and Functionally Distinct"

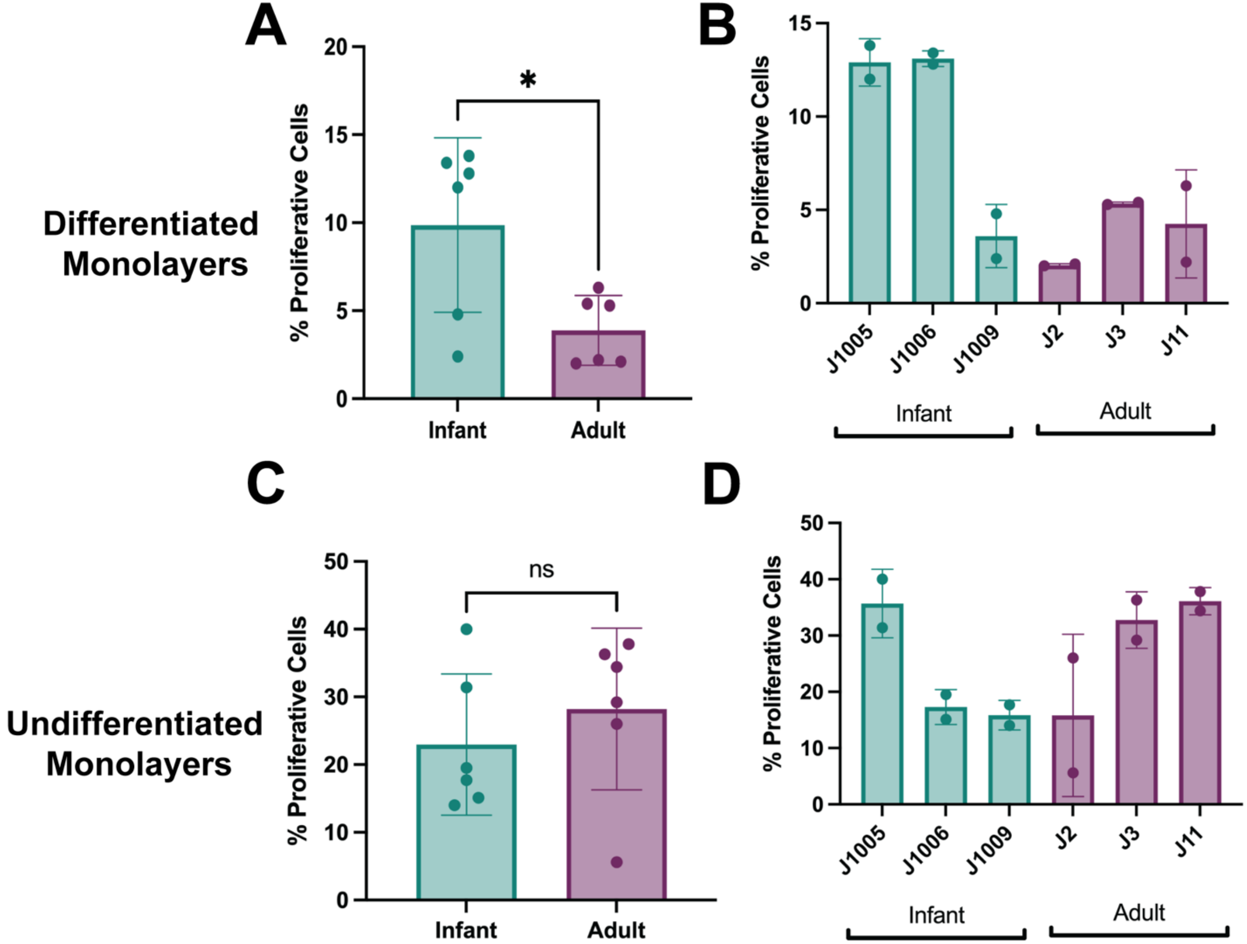


**Supplemental Figure 2: There are few proliferating cells in differentiated infant and adult HIE monolayers on transwells**

Percentage of EdU-positive cells quantified by flow cytometry in differentiated (A) and undifferentiated (C) HIE monolayers. B&D: Percentage of EdU-positive cells quantified by flow cytometry in individual lines. Data represents mean ± standard deviation (SD) from two independent experiments, with each experiment including the three infant and three adult HIE lines.
