## Supplemental Figure 3 for "Infant and Adult Human Intestinal Enteroids are Morphologically and Functionally Distinct"

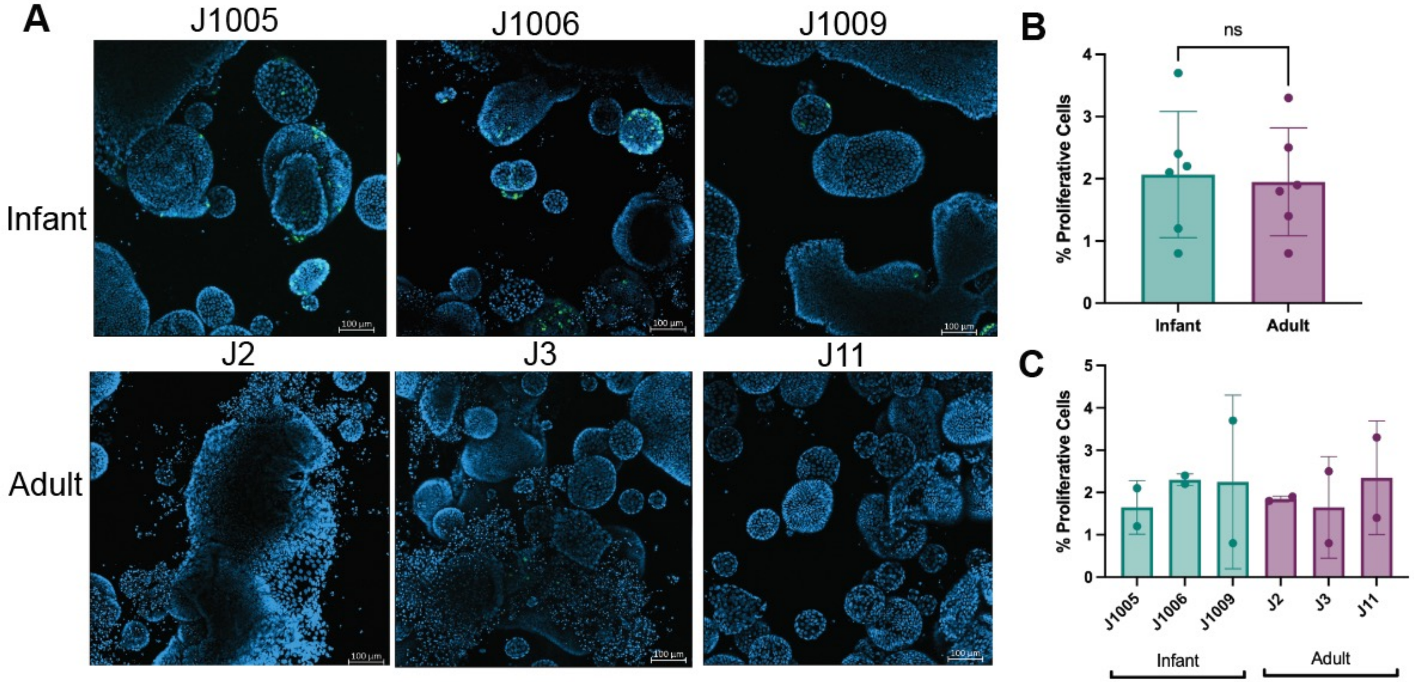


**Supplemental Figure 3: There are few proliferating cells in differentiated infant and adult 3D HIEs**

A: Representative confocal 3D reconstruction images after 24h 5-ethynyl-2′-deoxyuridine (EdU) incorporation in differentiated infant and adult HIEs. B: Percentage of EdU-positive cells quantified by flow cytometry. C: Percentage of EdU-positive cells quantified by flow cytometry in individual lines. Data represents mean ± SD from two independent experiments, with each experiment including the three infant and three adult HIE lines. The *p*-values were calculated by student's t-test.
