## Supplemental Table 1 for "Infant and Adult Human Intestinal Enteroids are Morphologically and Functionally Distinct"

| GeneID | GeneSymbol | GeneBiotype | logFC | PValue | FDR |
|---|---|---|---|---|---|
| ENSG00000104067 | TJP1 | Protein coding | 0.000 | 0.999 | 1.000 |
| ENSG00000119139 | TJP2 | Protein coding | -0.292 | 0.511 | 0.780 |
| ENSG00000105289 | TJP3 | Protein coding | -0.500 | 0.046 | 0.186 |
| ENSG00000197822 | OCLN | Protein coding | 0.139 | 0.578 | 0.827 |
| ENSG00000163347 | CLDN1 | Protein coding | -1.966 | 0.000 | 0.000 |
| ENSG00000165376 | CLDN2 | Protein coding | 4.368 | 0.000 | 0.000 |
| ENSG00000165215 | CLDN3 | Protein coding | 0.440 | 0.129 | 0.370 |
| ENSG00000189143 | CLDN4 | Protein coding | -0.200 | 0.455 | 0.740 |
| ENSG00000181885 | CLDN7 | Protein coding | 0.122 | 0.649 | 0.862 |
| ENSG00000157224 | CLDN12 | Protein coding | -0.544 | 0.031 | 0.140 |
| ENSG00000106404 | CLDN15 | Protein coding | 0.795 | 0.002 | 0.015 |
| ENSG00000113946 | CLDN16 | Protein coding | -1.918 | 0.003 | 0.023 |
| ENSG00000066405 | CLDN18 | Protein coding | -3.358 | 0.000 | 0.000 |
| ENSG00000253958 | CLDN23 | Protein coding | -1.437 | 0.000 | 0.000 |

**Supplemental table 1**

RNA Sequencing data on tight junction proteins expressed in infant over adult HIEs.
