## Supplemental Figure 4 for "Infant and Adult Human Intestinal Enteroids are Morphologically and Functionally Distinct"

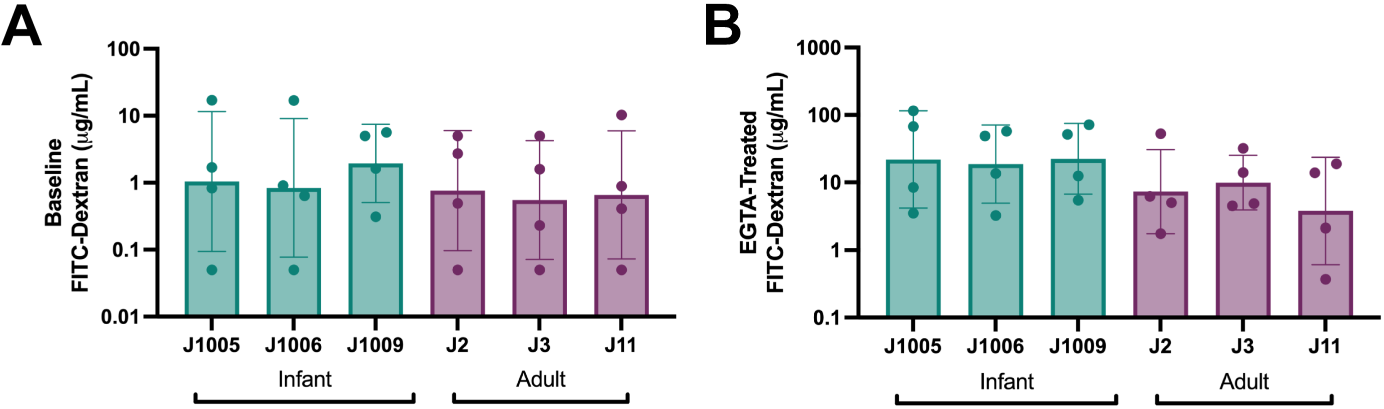


**Supplemental Figure 4: FITC-Dextran concentration in response to EGTA treatment by line**

Concentration of 4kDA FITC-Dextran in each HIE line at (A) baseline and (B) after EGTA treatment. Data represent mean ± SD from four independent experiments, with each experiment including the three infant and three adult HIE lines.
