## Supplemental Figure 5 for "Infant and Adult Human Intestinal Enteroids are Morphologically and Functionally Distinct"

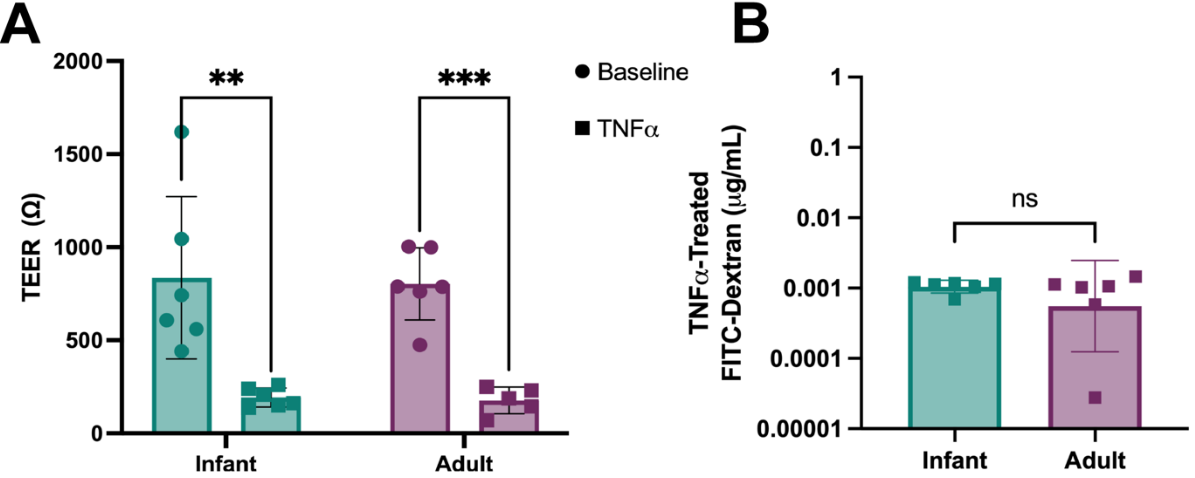


**Supplemental Figure 5: TNFα causes changes in TEER in both infant and adult HIEs.**
A: TEER values of HIEs at baseline and after TNFα treatment. B: Concentration of 4kDA FITC-Dextran after TNFα treatment. Data represent mean ± SD from two independent experiments, with each experiment including the three infant and three adult HIE lines. The *p*-values were calculated by student's t-test, and the asterisks (**) and (***) represent $p < 0.01$, and $p < 0.001$ respectively
