## Supplemental Figure 7 for "Infant and Adult Human Intestinal Enteroids are Morphologically and Functionally Distinct"

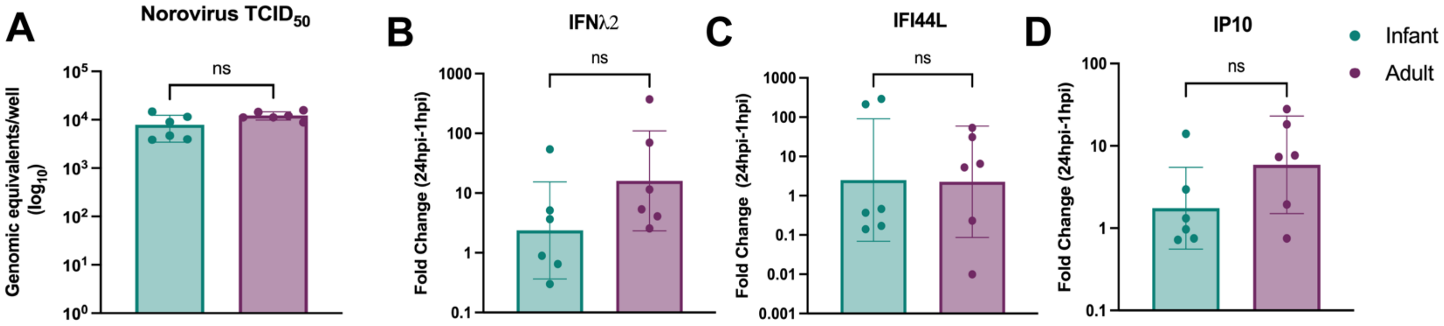


**Supplemental Figure 7: HuNoV tissue culture infectious dose 50 ($TCID_{50}$) and innate immune response to HuNoV infection**

A: HuNoV (GII.4) tissue culture infectious dose 50 ($TCID_{50}$) in HIEs at 24hpi. Quantification of (B) IFNλ2, (C) IFI44L, and (D) IP10 transcripts in response to infection with 100 TCID50s of HuNoV. Data represent mean ± SD from two independent experiments, with each experiment including the three infant and three adult HIE lines. The *p*-values were calculated by student's t-test.
