## Supplemental Figure 9 for "Infant and Adult Human Intestinal Enteroids are Morphologically and Functionally Distinct"

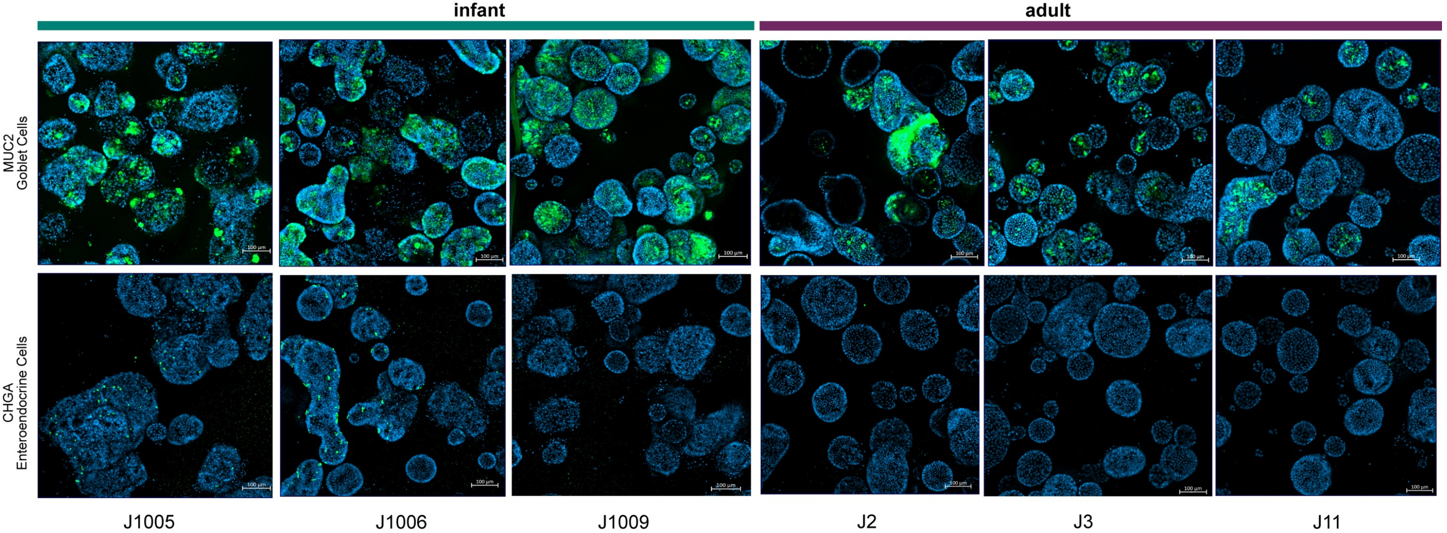


**Supplemental Figure 9: Cell type composition of varies between differentiated 3D infant and adult HIEs**

A: Representative confocal 3D reconstruction images of differentiated cell types in infant and adult HIEs. Top panel: goblet cells (Muc2, green), and bottom panel: enteroendocrine cells (ChgA, green). Nuclei are stained with DAPI (blue), Scale bar = 100 µm.
