## Supplemental Figure 10 for "Infant and Adult Human Intestinal Enteroids are Morphologically and Functionally Distinct"

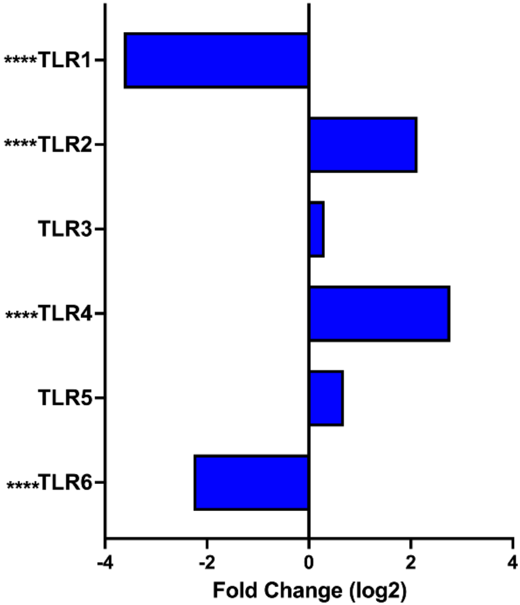


**Supplemental Figure 10: Expression of some toll-like receptors (TLRs) are significantly different in infant HIEs**

Expression profile of RNA-seq data for TLR genes. Data represents mean values and are expressed as Log2 fold change, the asterisk (****) represents $p < 0.0001$.
