## Supplemental Figure 11 for "Infant and Adult Human Intestinal Enteroids are Morphologically and Functionally Distinct"

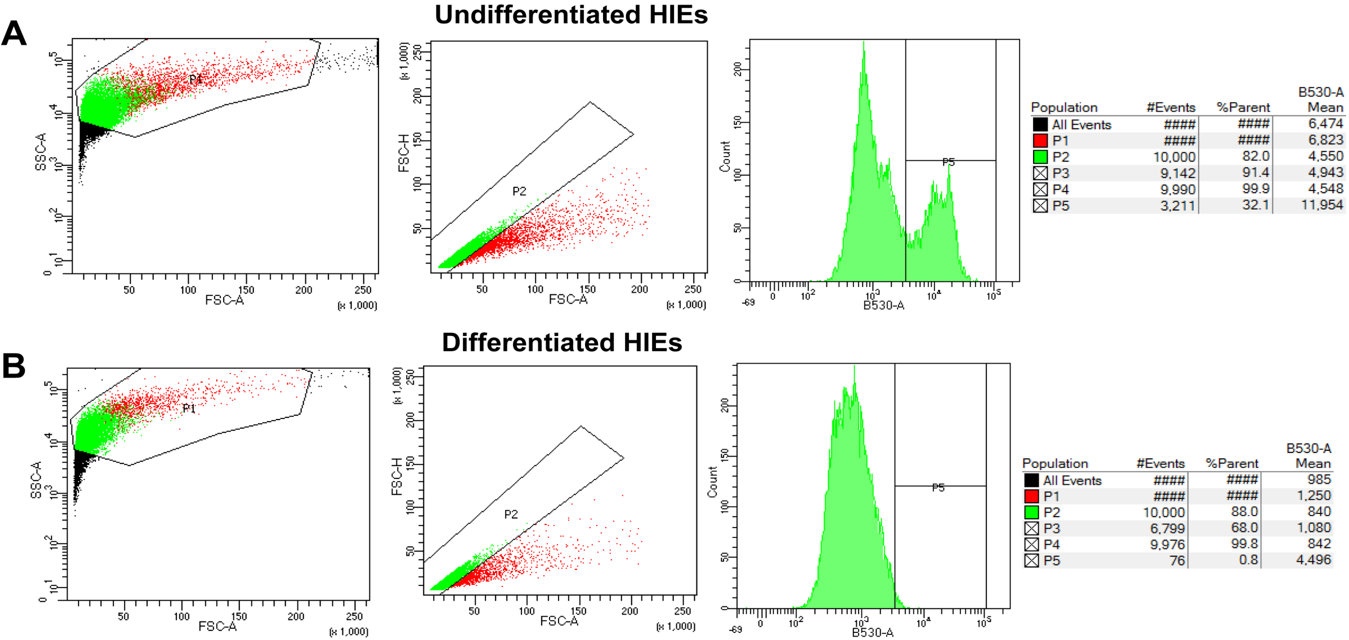


**Supplemental Figure 11:** Representative EdU gating strategy for undifferentiated (A) and differentiated (B) HIEs
